# The vaginal microbiota, symptoms, and local immune correlates in transmasculine individuals using sustained testosterone therapy

**DOI:** 10.1101/2025.03.14.643255

**Authors:** Bern Monari, Hannah Wilcox, Priscilla Haywood, Pawel Gajer, Jorge Rojas-Vargas, David Zuanazzi, Lindsay Rutt, Ainslie Shouldice, Reeya Parmar, L. Elaine Waetjen, Yonah Krakowsky, Emery Potter, Jessica L. Prodger, Jacques Ravel

## Abstract

Transmasculine individuals (assigned female at birth, masculine gender identity, TM) may use gender-affirming testosterone therapy, and some TM report adverse genital symptoms during treatment. In cis women, the vaginal microbiota is a key determinant of reproductive and sexual health outcomes; *Lactobacillus*-dominant communities are considered optimal, while more even, diverse, *Lactobacillus*-depleted microbiota are considered non-optimal. Prior studies suggest *Lactobacillus* deficiency in TM vaginal microbiota, but associations with symptoms and immune markers remain unclear. We launched the TransBiota study to characterize the TM vaginal microbiota, soluble mediators of local inflammation (SMI), and self-reported symptoms over three weeks. Fewer than 10% of TM possessed *Lactobacillus*-dominant microbiota, and most exhibited more diverse, *Lactobacillus-*depleted microbiota. We identified 11 vaginal microbiota community state types (tmCSTs), with *Lactobacillus*-dominant tmCSTs unexpectedly linked to abnormal odor and elevated IL-1α. These findings indicate that *Lactobacillus* dominance may no longer be an optimal state for TM during gender-affirming testosterone therapy and change in clinical management is needed.

## INTRODUCTION

Recent estimates from UCLA’s School of Law Williams Institute reveal that approximately 0.5% of US adults, more than 1.3 million individuals, identify as transgender or gender diverse (TGD), signifying that their gender identity does not align with their sex assigned at birth^1-4^. While gender is traditionally viewed as binary (male or female), TGD individuals may express a broad spectrum of gender modalities, including binary gender, absence of gender identity, identities outside the male/female framework, or multiple gender identities^1,5^. Transmasculine (TM) individuals were assigned female at birth but experience a male or masculine gender identity^5^. Conversely, cisgender (cis) individuals experience alignment between their sex assigned at birth and gender identity^5^. TGD individuals may pursue various pathways to align their external gender presentation with their internal sense of identity, such as social and behavioral changes, hormone therapy, and surgical interventions^6^. Although not all TGD individuals opt for medical transition, 78% of respondents to the 2015 United States Transgender Survey (USTS) expressed a desire for or use of hormone therapy, a proportion that rises to 95% among those with binary gender identities^1^. Early insights from the 2022 USTS further indicate that over 90% of TGD individuals receiving hormone therapy report improved life satisfaction^7^. In TM individuals, gender-affirming testosterone therapy is commonly utilized and administered through injections, gels, or patches^8-12^. Sustained testosterone use (>1 year) in TM has been shown to elevate serum testosterone levels to those typically seen in cis males (CM), while serum estrogen levels decrease to the low levels seen during menstruation in cis women of reproductive age (rCF)^13-15^. TM individuals undergoing sustained testosterone therapy frequently report adverse genital symptoms, such as vaginal atrophy, dryness, irritation, non-menstrual bleeding, and pain during sex^9^. Fewer than 2% of TM individuals have undergone genital surgeries, including metoidioplasty, phalloplasty, or vaginectomy^1,16^. Consequently, many TM retain a vagina exposed to high testosterone and low estrogen levels, often accompanied by vaginal symptoms.

In rCF, the vaginal microbiota plays a crucial role in reproductive and sexual health outcomes. Unlike at other body sites, the rCF vaginal microbiota is commonly dominated by a single species of *Lactobacillus*. Most *Lactobacillus*-dominated vaginal microbiota are considered “optimal” in cis women, based on observational studies that linked *Lactobacillus* dominance to a reduced risk for HIV and sexually transmitted infection (STI) acquisition, although *L. iners*-dominated vaginal microbiota may provide less protection compared to other *Lactobacillus* spp.-dominated microbiota^17-25^. Non-*Lactobacillus*-dominated vaginal microbiota typically consist of facultative and obligate anaerobes at relatively even abundances, including *Gardnerella*, *Fannyhessea*, *Prevotella*, *Sneathia*, *Dialister*, *Megasphaera*, and *Mobiluncus* species^21,26-29^. These more diverse and even microbiota have been associated with adverse outcomes in rCF, such as preterm birth^30-32^, bacterial vaginosis^25,33,34^, STIs including chlamydia and gonorrhea^24,34-37^, and HIV acquisition^27,38-40^. These associations have not yet been explored in TM. In rCF, genital tract microbiota can vary widely among individuals, with some experiencing rapid and frequent changes while others maintain stable microbiota over time^33,41-43^. To date, no studies have assessed the stability of the vaginal microbiota in TM individuals or its potential implications for health outcomes.

Seven broad community state types (CSTs) have been identified in a large study of the vaginal microbiota in rCF (>13,000 samples)^29^. Four CSTs are defined by the dominance of a single *Lactobacillus* species: CST I is dominated by *L. crispatus*, CST II by *L. gasseri*, CST III by *L. iners*, and CST V by *L. jensenii*^29^. CSTs I and III are further divided into CST I-A, CST I-B, CST III-A, and CST III-B, where communities assigned to the A subtype have higher relative abundance of the dominant taxa than communities assigned to the B subtype.^29^ The remaining three CSTs, CSTs IV-A, IV-B, and IV-C, are characterized by a low relative abundance of lactobacilli and a diverse set of strict and facultative anaerobic bacteria^29^. CST IV-A is characterized by a higher relative abundance of *Candidatus* Lachnocurva vaginae (formerly BVAB1) and moderate proportions of *Gardnerella vaginalis* and *Fannyhessea vaginae*. CST IV-B has a higher relative abundance of *G. vaginalis* with moderate levels of *Ca.* L. vaginae and *F. vaginae*^29^. CST IV-C is characterized by a low relative abundance of *Lactobacillus* spp., *G. vaginalis, F. vaginae,* or *Ca.* L. vaginae, and comprises five sub-CSTs with varying dominant bacterial taxa: CST IV-C1 has a higher relative abundance of *Streptococcus spp.*, CST IV-C2 of *Enterococcus* spp., CST IV-C3 of *Bifidobacterium* spp., and CST IV-C4 of *Staphylococcus* spp., and CST IV-C0 of *Prevotella* spp.

A prior cross-sectional study of 28 TM individuals accessing sustained testosterone therapy for at least one year found that most participants had vaginal microbiota devoid of *Lactobacillus* spp^44^. Instead, more diverse microbiota were observed, often resembling CST IV-C^44^. However, the study did not evaluate the microbiota in relation to symptoms or local inflammation, which are key factors in genital health. To address these gaps, we initiated the TransBiota study to comprehensively characterize the composition and structure of the vaginal microbiota, soluble mediators of local inflammation (SMI), and self-reported symptoms and behaviors in transgender individuals over three weeks. Our findings reveal that less than 10% of sampled TM exhibit *Lactobacillus*-dominant communities, with unique and diverse microbiota reminiscent of postmenopausal women. Interestingly, nearly half of TM individuals report genital symptoms, and *Lactobacillus-*dominance was associated with increased reports of abnormal or unpleasant odor, as well as elevated IL-1α concentration, a stark contrast to what has been observed in rCF^38,45-48^. These findings suggest that *Lactobacillus* dominance may no longer represent an optimal state for transmasculine individuals taking testosterone.

## RESULTS

### Study Participants Characteristics

We analyzed the vaginal microbiota, vaginal pH, and local SMI sampled at three weekly time points (n=208) in 85 TM individuals enrolled in the TransBiota study^49^, all of whom had been undergoing testosterone therapy for at least one year. Demographic characteristics of the TransBiota TM cohort are detailed in Table 1. Over half of the participants (56.6%) reported binary identity, while 37.6% reported a non-binary, genderqueer, agender, or similar identities. Two participants (2%) identified with indigenous or culturally specific genders, and three participants reported alternative gender identities within the transmasculine spectrum. The majority of participants identified as white and non-Hispanic (65.9%), with approximately a quarter (24.7%) identifying as multi-racial. The mean age of respondents was 29.8 years (range: 18-45), with the majority (57.6%) being 30 years old or younger. On average, participants reported taking testosterone for 5.1 years (range: 1.1-18.9 years). Table S1 outlines additional cohort-specific characteristics. Approximately 20% of participants reported undergoing gender-affirming surgeries involving the reproductive system, though individuals who had undergone vaginectomy were excluded from the present study. Reported surgeries included hysterectomy, oophorectomy, and metoidioplasty (clitoral release). The survey did not collect data on chest masculinization procedures.

**Table 1.**
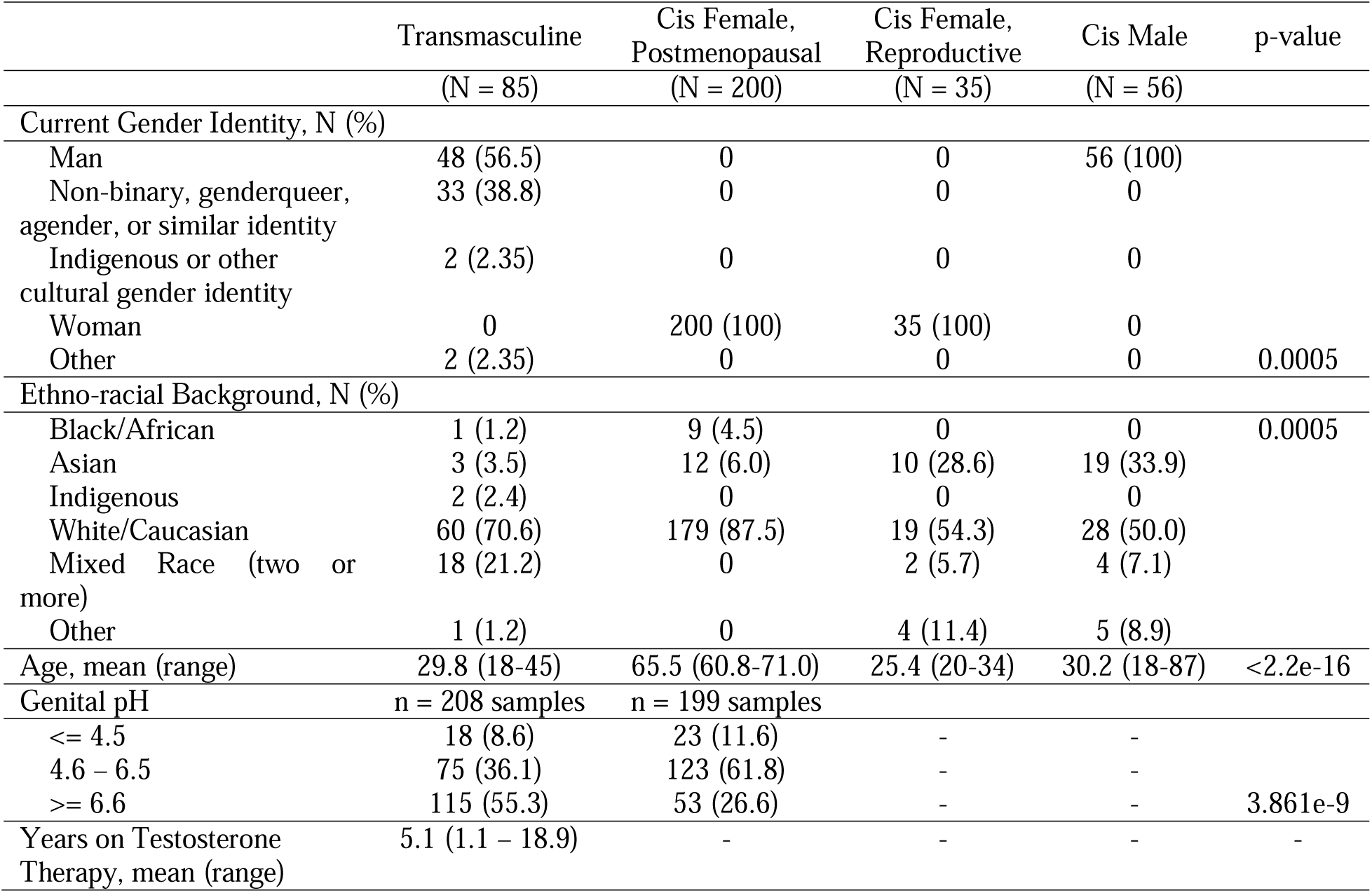
Demographic characteristics of the cohort included in the study. p-values are either Fisher’s exact test (categorical) or Kruskal-Wallis (continuous)

Regarding other health-related factors, three participants reported taking the anti-androgenic medication finasteride, often used to treat male-pattern baldness, while four participants utilized vaginal estradiol during the study period. One participant reported a recent gonorrhea diagnosis (within the past 30 days), but no participants indicated a recent chlamydia diagnosis. Only one participant reported douching in the prior 30 days. Notably, 43.5% of participants experienced at least one genital symptom in the week leading up to sampling, with the most commonly reported symptoms being vaginal dryness (23.5%) and pain during sex (12.9%) (Table S3).

We also recruited 35 rCF to characterize their vaginal microbiota and SMI, as well as 56 uncircumcised CM to characterize their penile microbiota (coronal sulcus). Additionally, we included 200 postmenopausal cisgender females (mCF) randomly selected from the Study of Women’s Health Across the Nation (SWAN) study, who had previously undergone vaginal microbiota characterization^50^. These rCF, CM and mCF groups served as comparator populations (Table 1). The comparator groups differed demographically from the TM cohort. Specifically, rCF, CM and mCF participants were more likely to identify as non-white, with an increased frequency of Asian ethno-racial backgrounds among CM and rCF. Age distribution also varied, with CM and mCF participants being older than both TM and rCF participants. In terms of vaginal pH, TM participants exhibited significantly higher pH levels compared to mCF participants, with 55.3% of TM having a vaginal pH over 6.6 compared to 26.6% of rCF (Table 1). However, pH data were not collected for the rCF and CM, limiting direct comparisons to these groups.

### The vaginal microbiota of TM accessing testosterone therapy does not resemble the CF vaginal microbiota nor the CM penile microbiota

We first visualized the relative abundances of the 50 most abundant bacterial taxa observed among TM, rCF, mCF, and uncircumcised CM individuals (Figure 1). *Lactobacillus* dominance was observed in fewer than 10% of TM individuals, compared to 82.9% of rCF, 26.5% of mCF, and only 3.5% of CM. In contrast, most TM individuals possessed a microbiota with relatively even relative abundances of strict and facultative anaerobes, including *Peptoniphilus*, *Anaerococcus*, *Dialister*, *Streptococcus*, and *Prevotella* species. One particularly notable TM individual harbored a microbiota dominated (≥95% relative abundance) by *F. vaginae* across all three sampling time points, a rare observation among rCF^51^. The penile microbiota of CM also demonstrated relatively even compositions, with moderately elevated relative abundances of *Prevotella* and *Corynebacterium* species. As expected, the rCF microbiota adhered to previously defined CSTs: 54.3% were assigned CST I (*L. crispatus*), 5.7% CST II (*L. gasseri*), 20% CST III (*L. iners*), and 5.7% CST V (*L. jensenii*). Five rCF samples (14.3%) were assigned to CST IV-B and included taxa such as *L. iners*, *G. vaginalis*, *F. vaginae*, and *Megasphaera* spp. In comparison, mCF samples were predominantly classified as CST IV-C (57.5%) or IV-B (7.5%), with sub-CST IV-C types distinguished by higher relative abundances of *Streptococcus* spp. (CST IV-C1) or *Bifidobacterium* spp. (CST IV-C3). CST IV-C1 was observed in both mCF and TM individuals, while CST IV-C3 was exclusive to mCF.

**Figure 1.**
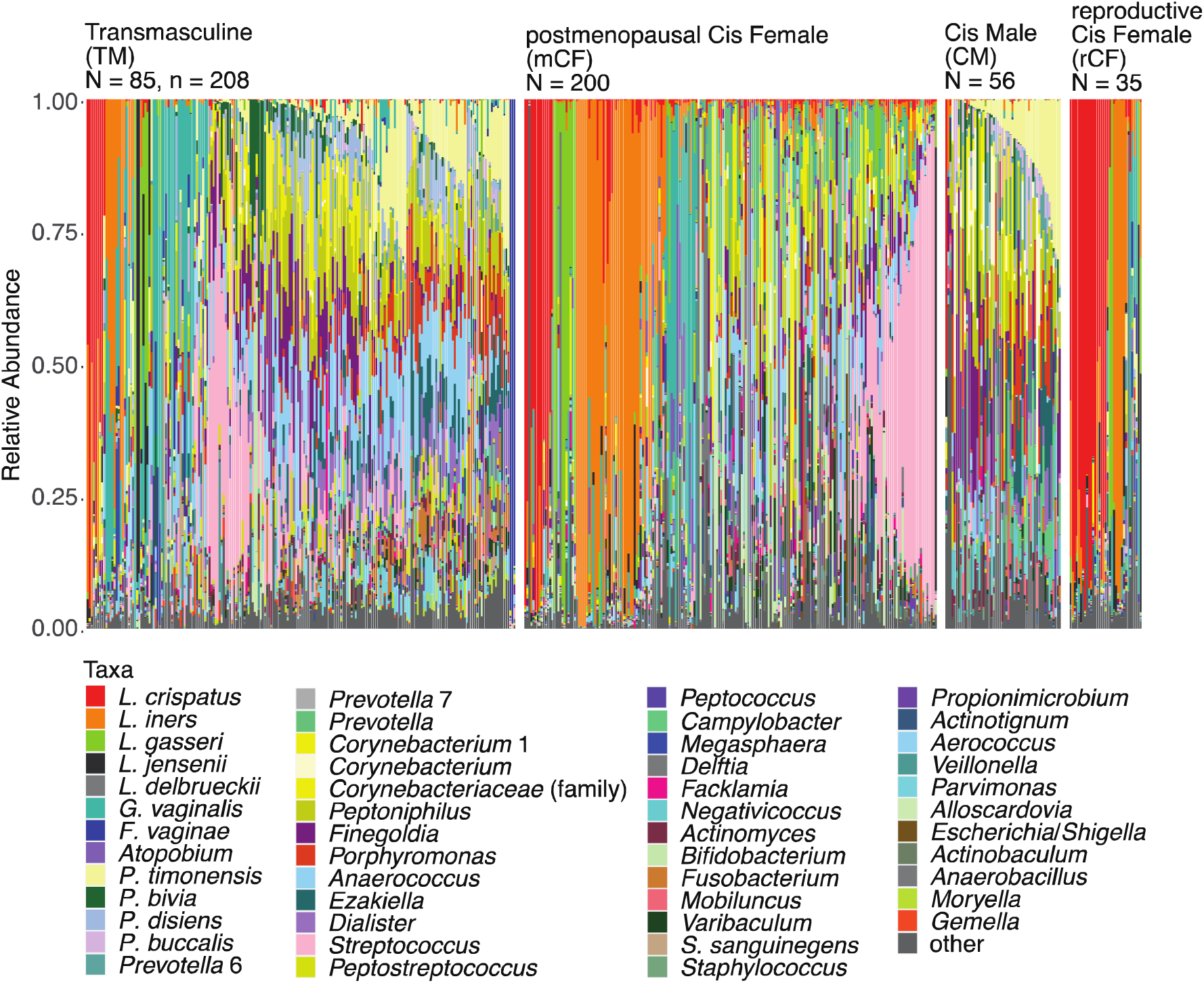
Bacterial taxonomic composition of the genital microbiota of TM, CM, rCF, and mCF. Within each study group, samples were sorted by CST or tmCST, relative abundance of *P. timonensis*, *Streptococcus* spp., *Corynebacterium* 1, and then *G. vaginalis*. The figure was slightly edited in Adobe Illustrator for aesthetics.

Principal coordinates analysis (PCoA) of Jensen-Shannon divergence measurements revealed key differences across the TM (n=208), mCF (n=200), rCF (n=35), and CM (n=56) groups. Figures 2A and 2B demonstrates that much of the variance when comparing all groups together was driven by *L. crispatus* and *L. iners,* taxa frequently found in the rCF vaginal microbiota but less common in other groups. Restricting the PCoA to TM, CM, and mCF microbiota (Figures 2C and 2D) revealed some overlap among these groups but with distinct clustering patterns. For instance, *L. iners* dominance in some mCF accounted for variation along both PC 1 and PC 2, masking finer differences among CST IV samples. To focus on the most diverse CST (IV-A, IV-B and IV-C), and better characterize TM vaginal microbiota, we limited the PCoA to include only samples from TM (n=183, 88%), mCF (n=131, 65%), and CM (n=54, 96.4%) that were assigned to these CSTs. This analysis revealed greater similarities between TM and mCF microbiota, with minimal overlap with CM microbiota. PC1 distinctly separated CM samples (Figures 2E and 2F), driven by taxa such as *Finegoldia*, *Peptoniphilus*, *Porphyromonas*, *Ezakiella*, and *P. timonensis*. A final PCoA comparing CST IV samples from TM (n=183, 88%) and mCF (n=131, 65%) participants (Figures 2G and 2H) revealed group-specific clustering. *Streptococcus, Prevotella* and *Anaerococcus* species were key drivers of mCF clustering patterns, while *P. timonensis* and *Porphyromonas* species were more characteristic of TM microbiota. These analyses demonstrate that the vaginal microbiota of most TM individuals accessing testosterone therapy is distinct from that of rCF and exhibits only limited similarities with mCF or CM microbiota. The specific composition and structure of the TM vaginal microbiota highlight its unique characteristics, underscoring the need for further investigation into its functional implications and potential health impacts.

**Figure 2.**
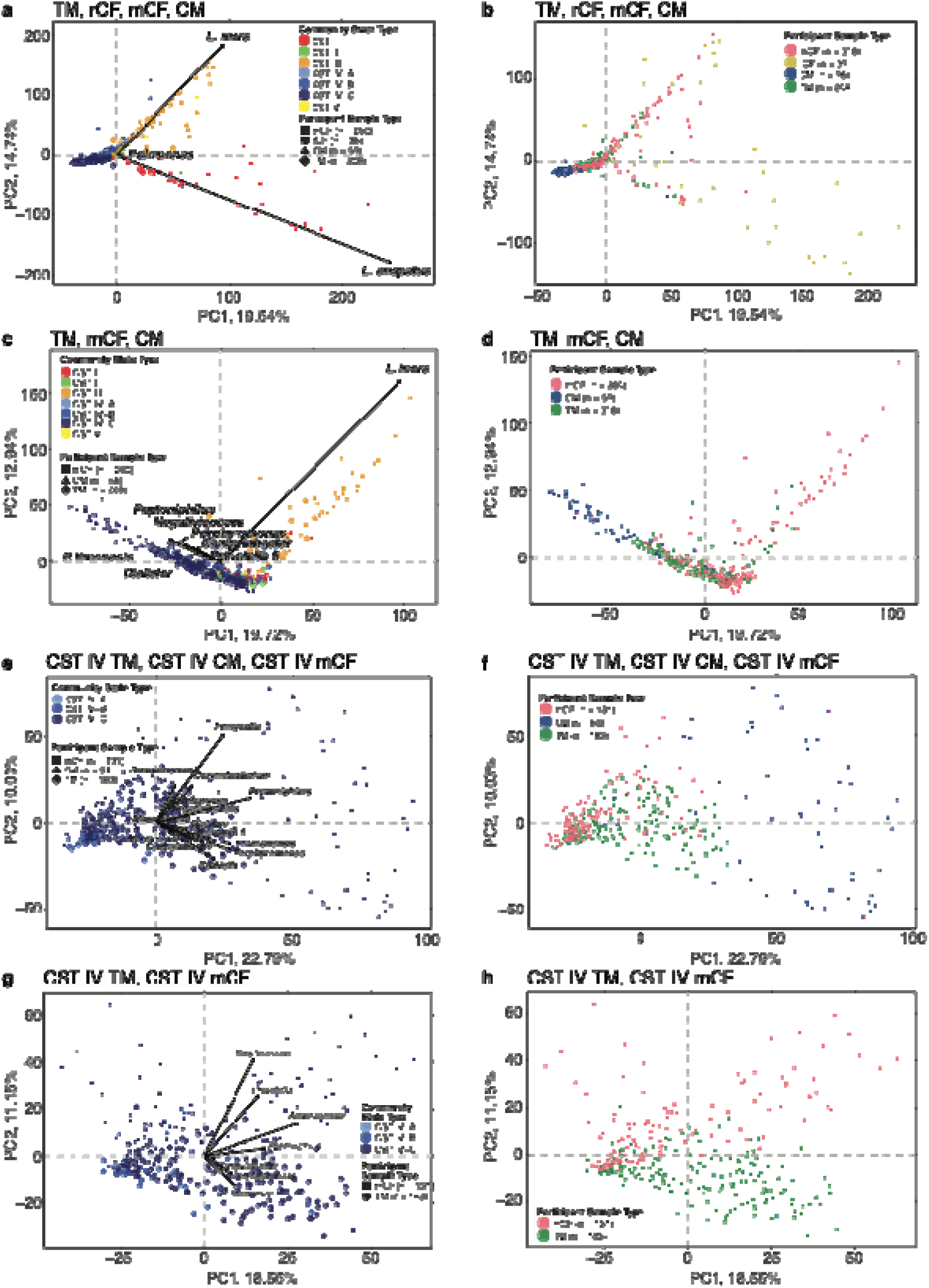
Principal Coordinates Analyses comparing the genital microbiota composition and structure of TM to rCF, CM, and mCF. Jensen-Shannon divergence matrix was calculated, and points were projected in 2D space. Significant drivers were determined using the *envfit* function in the BioDiversity R package. **a** and **b** all individuals from the four study groups. **c** and **d** all TM, all CM, and all mCF. **e** and **f** TM, mCF, and CM assigned to CST IV-A, IV-B, or IV-C. **g** and **h** transmasculine and postmenopausal samples assigned to CST IV-A, IV-B, and IV-C. The figure was slightly edited in Adobe Illustrator for aesthetics.

### Composition and Structure of the TM Vaginal Microbiota

Given that CSTs commonly observed in rCF did not sufficiently capture the variation observed in TM microbiota, we utilized PHATE (Potential of Heat-diffusion for Affinity-based Trajectory Embedding) to embed the microbiota data (n=208) in three-dimensional space. Initial embedding of TM microbiota samples isolated three samples, all from the same participant, and formed their own cluster. PHATE embedding of the remaining 205 samples was then clustered using hierarchical Density-Based Spatial Clustering of Applications with Noise (hDBSCAN), identifying ten distinct clusters, which we named transmasculine community state types (tmCSTs) (Figure 3A). The first five tmCSTs demonstrate some resemblance to CSTs observed in rCF and mCF: tmCST 1, 2 and 3 were characterized by elevated relative abundances of *Lactobacillus* species, with tmCST 1 (n=9 samples, N=5 participants) featuring high relative abundance of *L. crispatus*, tmCST 2 (n=14, N=9) of *L. iners*, and tmCST 3 (n=9, N=6) of *L. jensenii* or *L. gasseri*. Notably, tmCSTs 1 and 2 resemble CST I-B and CST III-B^29^, in which high relative abundance of *L. crispatus* and *L. iners* is accompanied by strict and facultative anaerobes, accounting for up to 40% relative abundance. tmCST 4 (n=26, N=15) was characterized by high relative abundance of *G. vaginalis*, similar to CST IV-B, which is often associated with bacterial vaginosis (BV) in rCF. tmCST 5 (n=21, N=14), dominated by *Streptococcus* spp., resembled CST IV-C1 microbiota commonly observed in mCF. The remaining tmCSTs 6 to 11 were distinct and not typically observed in CF populations. tmCSTs 6 and 8 were characterized by elevated relative abundance of a specific *Prevotella* spp., tmCST 6 (n=8, N=5) with *P. bivia* and tmCST 8 (n=16, N=11) with *P. timonensis.* Both clusters also exhibited *Streptococcus*, *Dialister*, *Anaerococcus*, and *Peptoniphilus* spp. in relatively even but lower relative abundances. tmCST 7 (n=52, N=31) displayed relatively even abundances of *Streptococcus*, *P. disiens*, *Anaerococcus*, *Peptoniphilus*, *Finegoldia*, and *Corynebacterium* 1, with lower relative abundances of *P. timonensis*, *P. bivia*, and *Dialister*. tmCST 9 (n=29, N=19) was characterized by even relative abundances of *P. timonensis*, *P. disiens*, *Anaerococcus*, *Peptoniphilus*, *Porphyromonas*, and *Ezakiella,* with a relatively low relative abundance of *Dialister*. tmCST 10 (n=21, N=16) was similar to tmCST 9 but had decreased *Anaerococcus* and elevated *Fusobacterium* relative abundances. Finally, all samples from participant TMI162 were uniquely classified as tmCST 11 (n=3, N=1), distinguished by 95% or greater relative abundance of *F. vaginae* at all three timepoints. These findings illustrate that, while a subset of TM microbiota resembled CSTs typically seen in CF populations, the majority (60.6% of samples) exhibited unique compositional and structural profiles not observed in cisgender individuals. This underscores the distinct microbial ecology within the vaginal environments of TM accessing testosterone therapy.

**Figure 3.**
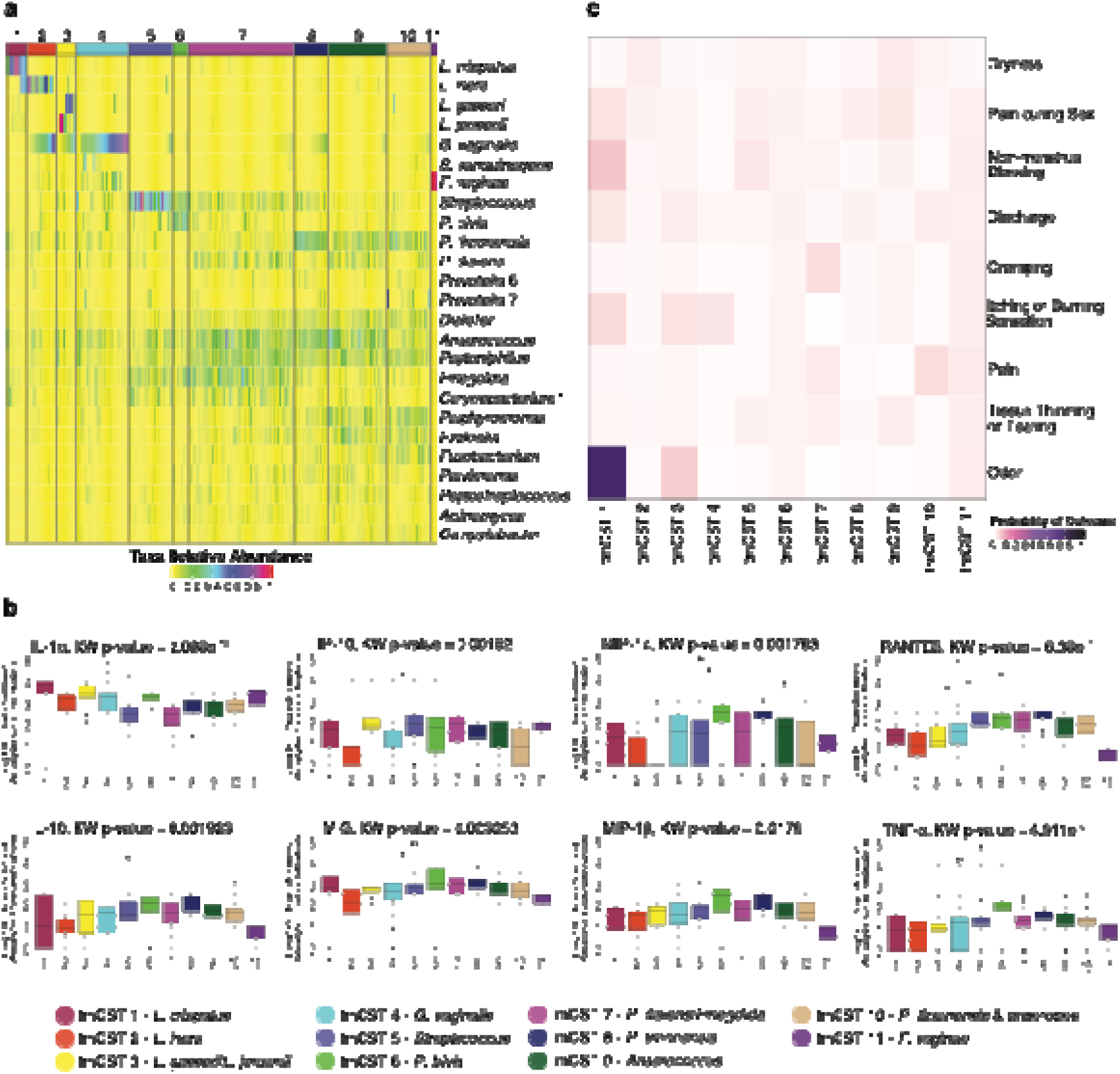
Community state types in the vaginal microbiota of transmasculine individuals (tmCST) **a** taxonomic composition of the 11 tmCSTs. **b** association between SMI and tmCSTs. Kruskal-Wallis was used to test for differences in distribution, where p <= 0.05 was deemed significant; Dunn’s Test with Bonferroni correction was implemented to identify pairwise differences between tmCSTs. * indicates p-value <= 0.05; ** indicates p-value <= 0.01; *** indicates p-value <= 0.001; **** indicates p-value <= 0.0001 **c** association between genital symptoms in the past 7 days and tmCSTs, determined using Bernoulli modeling with random effects correction for multiple samples from the same subject. The figure was slightly edited in Adobe Illustrator for aesthetics.

### Soluble markers of inflammation in the TM and rCF cohorts and associations with tmCSTs

We compared SMI concentrations between TM and rCF, the two cohorts for which SMI measurements were collected (Table S3). IL-8 was detected in all samples, and IL-6 was present in 98.1% of TM samples, but there were no significant differences in their concentrations between the two groups. IL-1α and IP-10 were lower in TM individuals, while the chemokines RANTES, MIP-1β, and MIP-1α, and the pro-inflammatory cytokine TNF-α, were significantly elevated in TM individuals compared to rCF. Additionally, the anti-inflammatory cytokine IL-10 was elevated in TM. Overall, TM individuals exhibited higher concentrations of most chemokines, TNF-α, and IL-10 compared to rCF, but lower levels of IL-1α and IP-10.

We next assessed the relationship between SMI concentrations and the composition and structure of the TM vaginal microbiota across the tmCSTs (Figure 3B). Analysis of log_10_-transformed concentrations revealed notable differences among tmCSTs. IL-1α concentrations were elevated in *Lactobacillus*-dominated tmCSTs 1 and 3 (*L. crispatus*, *L. gasseri*, and *L. jensenii*) compared to tmCSTs 5 (*Streptococcus*) and 7 (characterized by a diverse microbiota including *Anaerococcus*, *Peptoniphilus*, *P. disiens*, and *Finegoldia*). Samples in tmCST 2 (*L. iners*-dominated) exhibited lower concentrations of IL-10, IP-10, and MIG compared to tmCSTs 7 and 8 (characterized by elevated *P. timonensis* relative abundance). TNF-α concentrations were significantly lower in tmCST 2 compared to tmCST 6 (elevated *P. bivia*) and tmCST 8 (elevated *P. timonensis*). RANTES concentrations were elevated in tmCST 8 compared to tmCST 1, 2, 4 (*G. vaginalis-*dominated), and 11 (*F. vaginae*-dominated).

These findings suggest that soluble markers of inflammation are strongly associated with the composition and structure of the TM vaginal microbiota, with specific tmCSTs exhibiting distinct SMI profiles. Elevated IL-1α in *Lactobacillus*-dominated tmCSTs and the association of TNF-α and RANTES with diverse and *Prevotella*-rich tmCSTs indicate that the microbiota’s composition has a measurable impact on local inflammation in TM individuals.

### Associations between tmCSTs and Self-Reported Symptoms

We assessed the association between tmCSTs and self-report of symptoms (Figure 3C). Individuals in tmCST 1 and tmCST 3 (*Lactobacillus* spp.-dominated) had higher probabilities of experiencing symptoms such as “not normal” or unpleasant odor, non-menstrual bleeding, itching or burning sensation, and discharge. In contrast, participants in tmCST 2 (*L. iners*-dominated) and tmCST 9 (characterized by *Anaerococcus*, *Peptoniphilus*, and *Porphyromonas*) had the highest probability of reporting vaginal dryness. Notably, there was a low probability of self-reported symptoms in tmCST 5, which was characterized by elevated relative abundance of *Streptococcus* spp. These associations suggest that, while *Lactobacillus*-dominated microbiota are typically associated with vaginal health in rCF, they may contribute to reported symptoms in TM individuals accessing testosterone.

### Microbiota compositional stability and tmCSTs

We next evaluated the compositional stability of the TM vaginal microbiota by analyzing weekly samples collected over three weeks (Figure 4). To assess stability, we calculated the probability of transitioning from one tmCST to another (Figure 4A) and determined Yue-Clayton theta indices for consecutive paired samples (week 1 and week 2, or week 2 and week 3), as shown in Table S4. More than half of the weekly samples from the same participant were assigned to the same tmCST, indicating relative compositional stability in TM vaginal microbiota over short timeframes. Samples assigned to tmCST 5 (characterized by elevated *Streptococcus*) and tmCST 10 (*P. timonensis* and anaerobe*-*enriched) were less likely to remain in the same tmCST in subsequent weeks, suggesting higher variability. Participants in tmCST 1 (*L. crispatus*-dominated) were more likely to remain in the same tmCST compared to tmCST 3 (*L. gasseri*- or *L. jensenii*-dominated), and individuals in tmCST 1 exhibited high Yue-Clayton theta indices, indicating a more stable microbiota in these participants. Participants in tmCST 10 had the lowest probability of staying in the same tmCST, yet paired samples assigned to tmCST 10 showed relatively high Yue-Clayton theta indices, suggesting substantial within-tmCST composition variation. Transitions between tmCSTs were visualized in Figure 4B, illustrating the relatively dynamic nature of TM vaginal microbiota composition. These analyses demonstrate that while TM microbiota exhibit relative stability, certain tmCSTs are more dynamic, with implications for their associations with symptoms and local inflammation.

**Figure 4.**
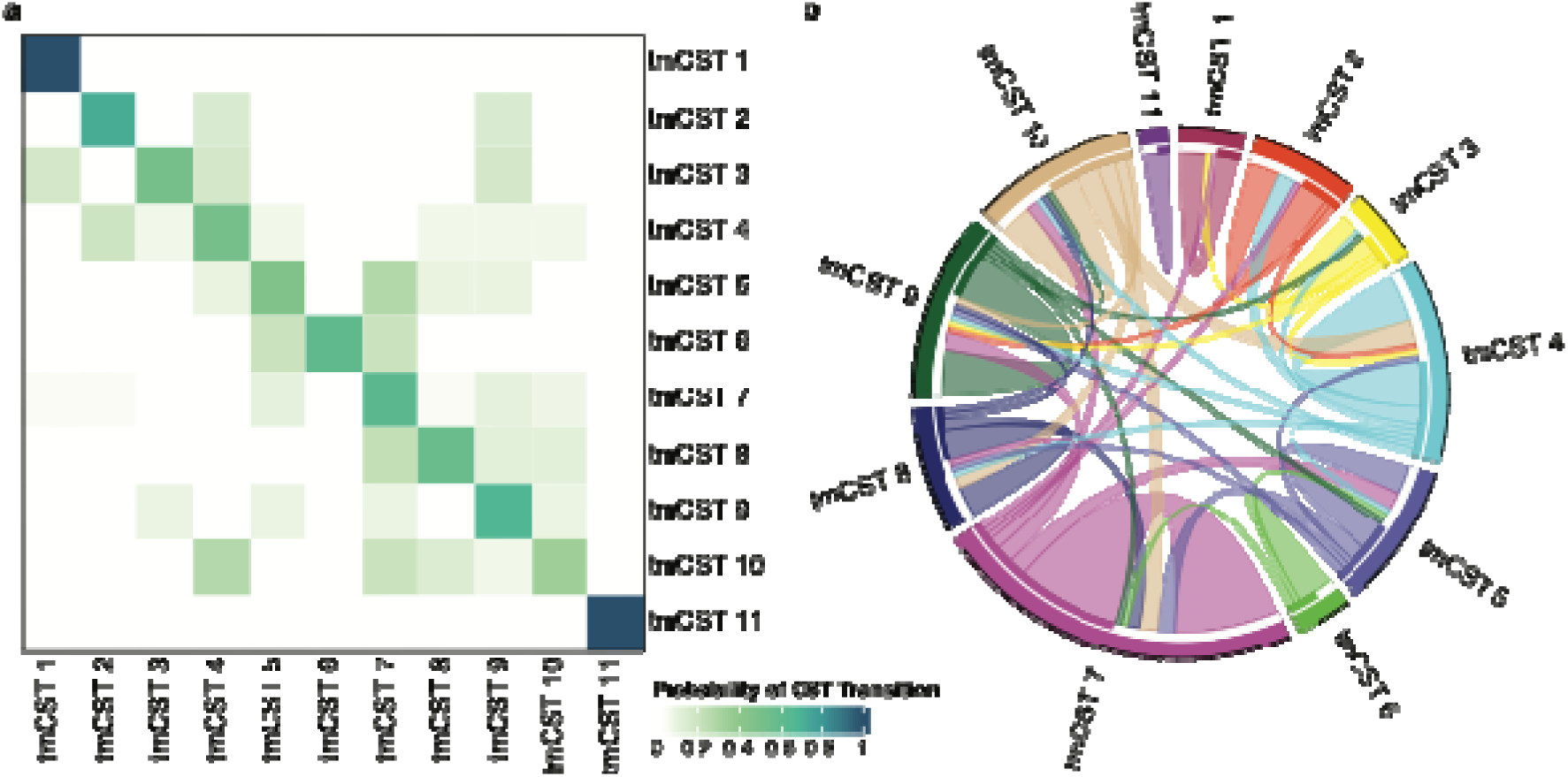
tmCST bacterial composition stability. **a** probability of a tmCST transitioning to another over the three weekly timepoints. **b** cord diagram representing the transitions between tmCSTs. The figure was slightly edited in Adobe Illustrator for aesthetics.

### Associations between bacterial relative abundances, symptoms, and SMI in TM

Hierarchical clustering of Spearman correlations between the top 50 bacterial taxa relative abundances in TM cohort samples revealed 10 distinct groups of bacterial taxa with positively correlated relative abundances, termed bacterial Taxa Clusters (TC) (Figure 5A). TC 1, 2, 3, and 4 were positively correlated with one another, as were TC 8 and 9, but these groups showed distinct SMI correlation profiles (Figure 5B). TC 10 contained *L. crispatus*, *L. gasseri*, and *L. jensenii*, commonly observed and considered optimal in rCF. These bacterial taxa were associated with low concentrations of SMI, except IL-1α and IP-10. TC 6 included *L. iners* alongside *Bifidobacterium*, *Megasphaera*, W5053, *Murdochiella*, and *Fastidiosipila*, bacterial taxa often observed in CST IV-C. This grouping suggests that *L. iners* may share characteristics with taxa associated with diverse microbiota in TM. TC 5 comprised taxa typical of CST IV in rCF (*Aerococcus*, *Sneathia sanguinengens*, *F. vaginae*, *Ureaplasma*, and *G. vaginalis*). These taxa were positively correlated with IL-1α, a marker of local inflammation. Several taxa were associated with self-reported symptoms (Figure 5C). For instance, *L. crispatus*, *L. gasseri*, and *L. jensenii* and *Corynebacterium* were positively associated with reports of abnormal or unpleasant odor. *Olsonella* (not in top 50 most abundant bacterial taxa) was positively associated with vaginal pain, while *P. bivia* was negatively associated with pain during sex. Taxa within TC 1 (*Peptoniphilus*, *Anaerococcus*, and *Actinomyces*), TC 2 (*P. disiens*), TC 4 (*Dialister*, *Campylobacter*, and *P. timonensis*), TC 8 (*Howardella*, *Finegoldia*, and *Streptococcus*), and TC 9 (*Facklamia* and *Corynebacterium* 1) were negatively associated with itching or burning sensation.

**Figure 5.**
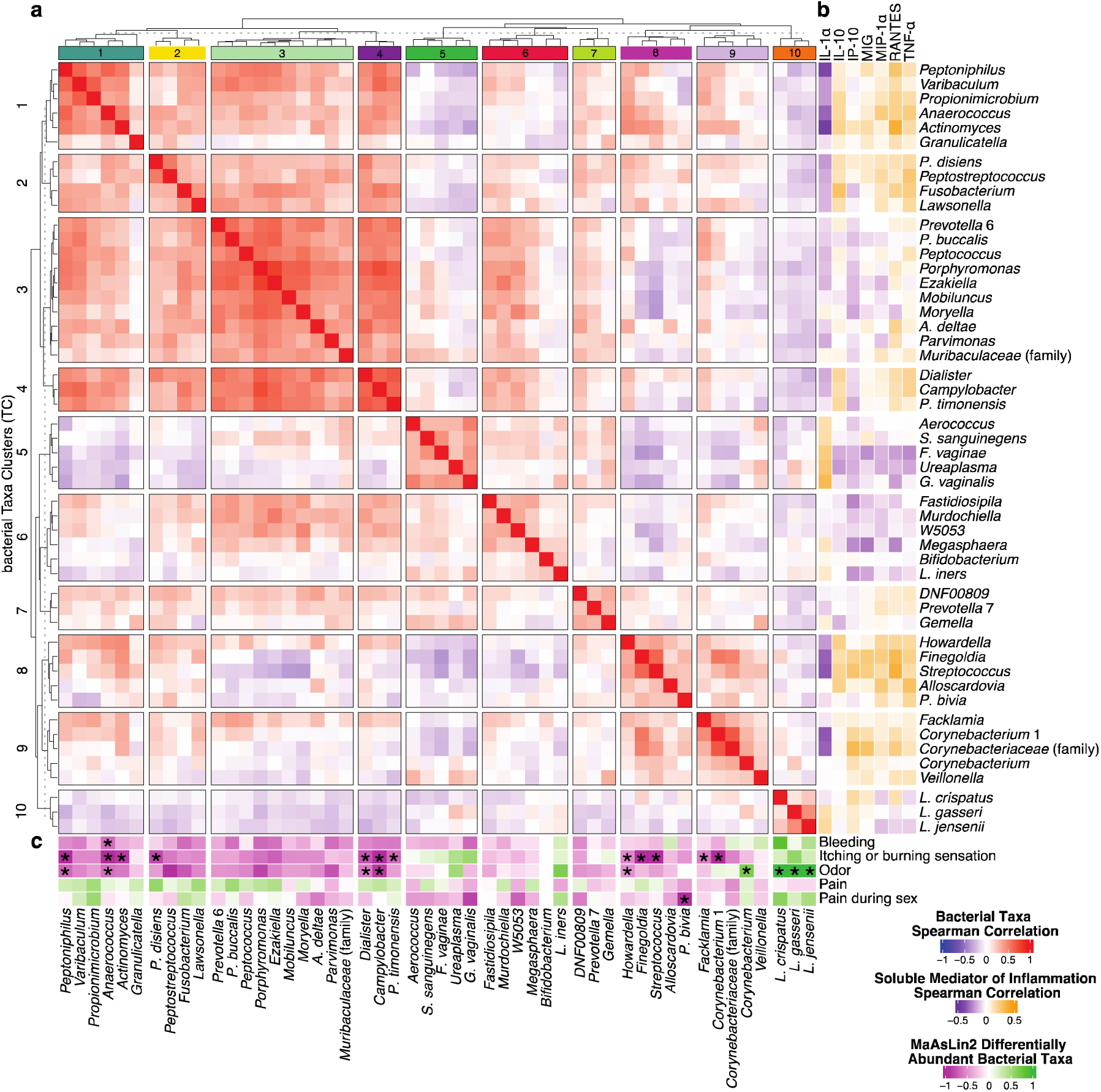
Correlations between the 50 most abundant bacterial taxa and soluble mediators of inflammation. **a** Spearman correlation analysis reveals 10 bacterial Taxa Clusters (TC) of positively correlated bacterial taxa. **b** Spearman correlations between 50 most abundant bacterial taxa and IL-1α, IL-10, IP-10, MIG, MIP-1α, RANTES, and TNF-α. **c** Differentially abundant bacterial taxa between individuals reporting and not reporting symptoms were identified using MaAsLin2, with significantly differentially abundant bacterial taxa denoted by an asterisk (*). The figure was slightly edited in Adobe Illustrator for aesthetics.

These findings highlight complex relationships between specific bacterial taxa, inflammatory markers, and self-reported symptoms in TM individuals. Traditional CSTs fail to capture the variation observed in the TM microbiota, underscoring the necessity of using tmCSTs and TCs to describe the unique microbial and inflammatory landscapes within this population.

### Local vaginal estradiol does not restore *Lactobacillus* dominance in TM

Four individuals in the study reported use of local vaginal estradiol, a treatment sometimes prescribed to TM taking testosterone. In mCF, local vaginal estradiol has been successfully used to improve vaginal epithelial thickness and, in some cases, restores *Lactobacillus* dominance in the vaginal microbiota^52-55^. The relative abundances of bacterial taxa observed at each timepoint for these individuals are shown in Figure 6. Although microbiota composition from samples of the same individual were more similar to each other than to those from other individuals, no specific bacterial taxa, tmCSTs, or TCs were associated with the use of local vaginal estradiol. These findings suggest that local vaginal estradiol may not have the same microbiota-restorative effects in TM individuals as observed in mCF.

**Figure 6.**
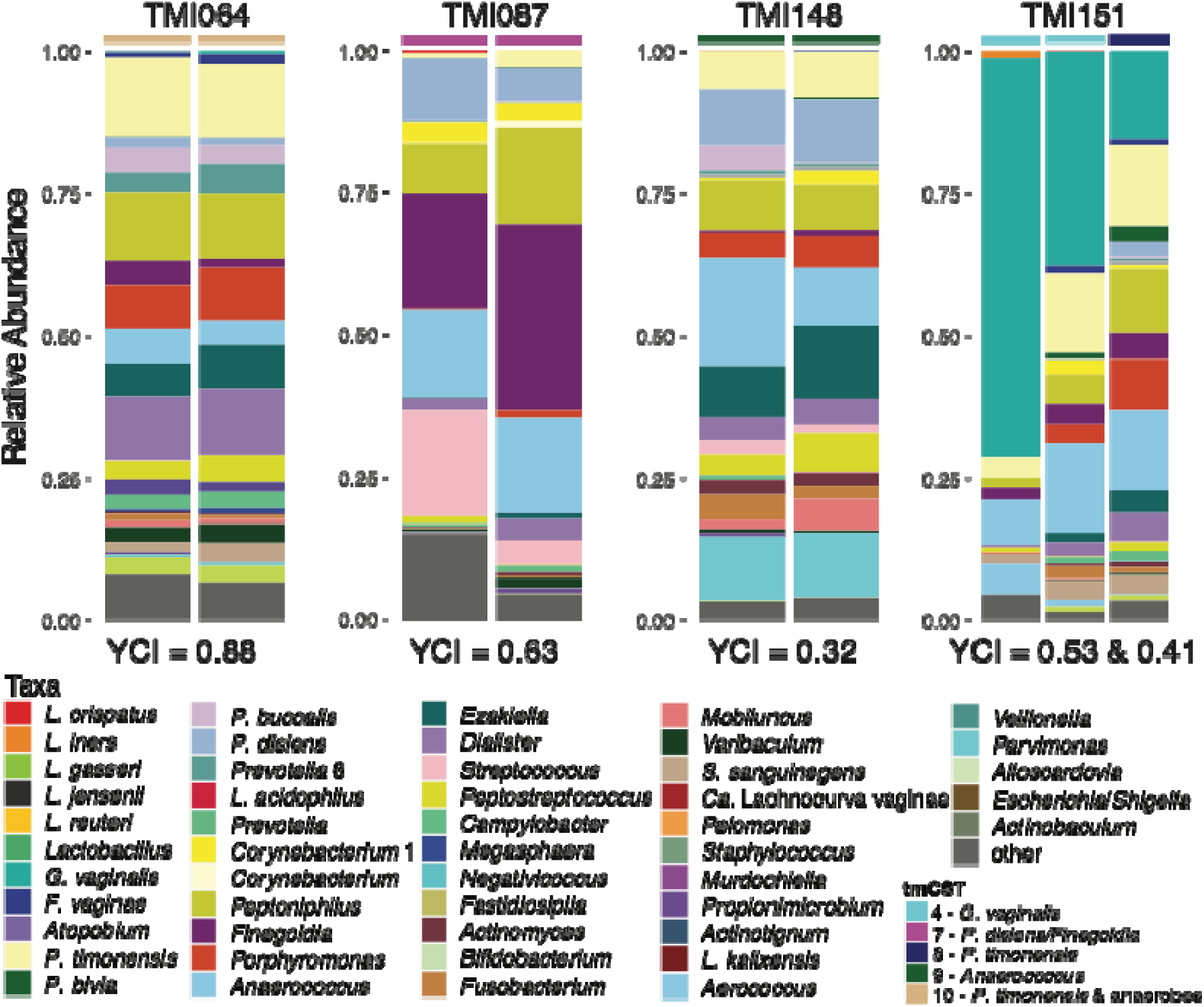
Vaginal microbiota composition in TM individuals using local estradiol. Yue-Clayton theta indices (YCI) were calculated using the designdist function from the R package *vegan* with the equation, “sum(x*y)/(sum(x^2^)+sum(y^2^)-sum(x*y))”. YCI can range from 0 to 1, with values closer to 0 indicating decreased similarity between pairs of samples and values closer to 1 indicating increased similarity between pairs of samples.

## Discussion

In the North American population, many individuals who identify as transgender or gender diverse are currently accessing or desiring gender-affirming hormone therapy^1,16^. Gynecological clinical care for TM individuals often follows guidelines designed for CF, which are based on the premise that *Lactobacillus*-dominant vaginal microbiota are optimal. However, it remains unclear whether vaginal microbiota deemed optimal for CF are also beneficial for TM individuals. Evidence-based guidelines tailored to the unique needs of TM are urgently needed to optimize their gynecological health. Critical gaps in knowledge remain regarding the composition and structure of the vaginal microbiota in TM individuals, as well as their association with symptoms, inflammation and adverse outcomes. This study sought to address these gaps by characterizing the vaginal microbiota of TM individuals who had been accessing testosterone therapy for over a year and examining associations with SMI and self-reported symptoms. We enrolled a broad cohort of 85 individuals to capture the diversity of vaginal microbiota in TM individuals on testosterone therapy.

In the present study, consistent with a previous report^44^, fewer than 10% of TM individuals exhibited a *Lactobacillus*-dominated microbiota. In contrast, rCF most frequently displayed this microbiota composition, where it is considered optimal, beneficial, and protective. Conversely, in rCF, CST IV-B vaginal microbiota, characterized by *G. vaginalis*, *F. vaginae*, and other strict and facultative anaerobes, are often associated with reports of symptoms^23,56^ and are considered non-optimal^30,57^. Interestingly, tmCST 4 in TM individuals resembled CST IV-B. Yet, a low proportion of TM individuals harboring these vaginal microbiota reported vaginal symptoms, suggesting that this profile may be more compatible with the testosteronized vagina. Conversely, an elevated relative abundance of *L. crispatus* and other *Lactobacillus* species in TM was associated with a higher probability of reporting abnormal or unpleasant genital odor, albeit these communities also contained low abundances of other bacterial anaerobes. These findings challenge the conventional understanding of *Lactobacillus* dominance as universally protective, suggesting that a high relative abundance of *Lactobacillus* spp. may not confer the same benefits in the testosteronized vaginal environment. The implications of these findings are significant, representing a potential paradigm shift. If reproduced in other cohorts, these results could alter the clinical and microbiological management of vaginal symptoms in TM individuals, particularly those related to odor which is believed to originate from microbial production of biogenic amines by vaginal bacteria. Further, this result does not support using Nugent scoring as a diagnostic for TM genital health, as it is likely to lead to unnecessary and potentially detrimental treatments.

Postmenopausal loss of estrogen is often associated with signs and symptoms such as vaginal atrophy, thinning, dryness, and dyspareunia, sometimes accompanied by urinary symptoms. Collectively, these symptoms are referred to as Genitourinary Syndrome of Menopause (GSM)^53,56,58^. Given the similar low serum levels of estrogens observed in TM individuals and the similarity of some of the symptoms they experience, we hypothesized that TM vaginal microbiota might resemble those of mCF. As previously reported^50^, *Lactobacillus* dominance is also less frequently observed in mCF. A high number of mCF were assigned to CST IV-C0 (31%), characterized by low but relatively even abundances of strict and facultative anaerobes such as *Anaerococcus*, *Peptoniphilus*, *Prevotella*, *Corynebacterium*, and *Propionimicrobium*. Additionally, 17% of mCF were assigned CST IV-C1, marked with elevated *Streptococcus* relative abundances, and 7.5% to CST IV-C3, characterized by elevated *Bifidobacterium*. While no TM individuals exhibited the *Bifidobacterium*-rich microbiota observed in CST IV-C3, the elevated abundance of *Streptococcus* species seen in CST IV-C1 was also observed in tmCST 5. In rCF, Group B *Streptococcus* has been associated with adverse obstetric outcomes, though it is often an asymptomatic colonizer of the gut and vaginal microbiota^59-61^. We postulate that tmCST 5-like microbiota may represent an optimal state for TM individuals, as it was associated with lower IL-1α concentrations and a reduced probability of self-reported symptoms. While this finding will require validation and reproduction in larger and more diverse cohorts, it represents an important step toward defining what constitutes optimal vaginal microbiota in TM individuals. Establishing a clearer understanding of the unique microbiota profiles associated with reduced inflammation and symptom burden in TM individuals could provide a foundation for evidence-based guidelines tailored to their gynecological health needs.

On the other end of the spectrum, uncircumcised CM were recruited as comparators based on the hypothesis that the microbiota associated with the less aerobic environment of the uncircumcised coronal sulcus, also featuring a stratified epithelium exposed to testosterone, might more closely resemble the testosteronized vaginal microbiota of TM individuals. The coronal sulcus microbiota of uncircumcised CM exhibited a diverse bacterial composition, including *Prevotella*, *Corynebacterium*, *Finegoldia*, *Ezakiella*, *Campylobacter*, *Anaerococcus*, *Porphyromonas*, and *Peptoniphilus* species, with particularly elevated relative abundance of *P. timonensis* and *P. disiens*. Indeed, a subset of TM individuals harbored penile-like microbiota (*e.g.,* tmCST 10). Despite the presence of *Prevotella* spp., which have been commonly associated with elevated pro-inflammatory cytokines and adverse outcomes in penile and vaginal environments^35,62-68^, these microbiota did not stimulate a strong pro-inflammatory response in TM individuals. Emerging evidence from *in vitro* studies suggests that *Prevotella* spp. alone do not inherently trigger significant pro-inflammatory responses but may facilitate the colonization of other anaerobic bacteria with pro-inflammatory properties, such as *G. vaginalis*^64^. This observation may indicate that penile-like microbiota in TM individuals potentially mitigating the adverse effects typically associated with *Prevotella* spp. in other environments. Interestingly, other tmCSTs also featured *Prevotella* species: tmCST 6 and 8 were characterized by elevated relative abundance (10-20%) of specific *Prevotella* spp. (*P. bivia* and *P. timonensis*, respectively), while *P. disiens* was consistently detected at low relative abundance in tmCST 7 and 9, often co-occurring with *P. timonensis* in tmCST 9. Additionally, *P. buccalis* was identified in multiple samples in the present and previous studies^69-71^. Among these *Prevotella*-enriched tmCSTs, immune activation was observed to be elevated but with varied significance. For instance, tmCST 8 exhibited the highest concentration of pro-inflammatory cytokines, supporting *in vitro* evidence that microbial context^64^, including the presence of co-occurring taxa, plays a critical role in determining the association between *Prevotella* spp., inflammation, and potentially adverse outcomes such as STI acquisition. However, it remains unknown whether the testosteronized vaginal epithelium responds to *Prevotella* spp. in the same manner as the estrogenized vaginal epithelium. These findings warrant further investigation *in vitro* to clarify the interplay between host hormonal environment, microbiota composition, and immune activation.

In the previous report on the testosterone-dominated vaginal microbiota^44^, two participants who reported using local vaginal estradiol exhibited a *Lactobacillus*-dominated microbiota, suggesting that local estradiol might elevate the relative abundances of *Lactobacillus* species in the vaginal microbiota of TM individuals. In rCF, estrogen stimulates glycogen accumulation in the superficial layers of the vaginal epithelium, a process that is thought to be critical for the establishment and maintenance of a *Lactobacillus*-dominant microbiota^72-74^. Glycogen serves as a key nutrient source for *Lactobacillus* species, which metabolize it into lactic acid^75-80^. The resulting acidic environment can inhibit the growth of anaerobic bacteria, thereby promoting *Lactobacillus* growth and colonization^20,57,76,81,82^. The importance of local estradiol in sustaining *Lactobacillus* spp. in the vagina has been demonstrated in mCF, where it improved vaginal epithelial integrity, restored *Lactobacillus* dominance, and in part, alleviated symptoms^50,52,53,56^. These observations have led to its occasional prescribed use by TM individuals to address similar symptoms. However, in the present study, the four TM individuals who reported using local estradiol did not exhibit high relative abundances of *Lactobacillus* spp. Instead, their microbiota displayed relatively even compositions of strict and facultative anaerobes, including *Anaerococcus*, *Finegoldia*, *P. timonensis*, and *Peptoniphilus*. This discrepancy suggests that concurrent estrogen and high testosterone levels in TM individuals may inhibit glycogen accumulation, which has been previously reported^72,83^ as necessary to sustain a *Lactobacillus*-dominant microbiota. Further *in vitro* and *in vivo* research is needed to elucidate the molecular mechanisms underlying the effects of local estradiol treatment on the testosteronized vaginal epithelium and microbiota. Studies exploring the interaction between local estradiol and systemic testosterone, alongside detailed characterization of epithelial and immune responses, could provide critical insights into tailoring therapies for TM individuals to optimize both microbiota composition and vaginal health. Ideally, these studies should be performed in the context of testosterone therapy initiation.

As our understanding of the unique physiological and microbiological changes in TM individuals grows, it becomes increasingly evident that current clinical practices often fail to address their specific needs. In clinical practice, TM individuals often receive care based on their sex assigned at birth, without consideration of the physiological and microbiological changes induced by masculinizing hormone therapy, hysterectomy, and oophorectomy, or other gender-affirming medical interventions. The findings from the present study underscore the need for tailored gynecological care for TM individuals, as *Lactobacillus-*dominated microbiota, typically considered optimal in rCF, were associated with non-optimal symptoms and local immune activations in TM. Conversely, microbiota lacking *Lactobacillus* spp. and comprising a wide array of different strict and facultative anaerobic bacteria were linked to reduced symptom reports and lower concentrations of immune mediators, suggesting that these states may not have the same adverse implications in TM. Further study is essential to delineate the effects of testosterone treatment on the cervicovaginal epithelium, including its impact on cervical mucus composition and the long-term shaping of the vaginal microbiota structure and functional capacity. Additionally, studies are needed to elucidate the molecular mechanisms and effectiveness of local estradiol treatment, vaginal moisturizers, or other interventions for managing vaginal dryness and related symptoms in TM individuals accessing testosterone treatment. Lastly, diagnostic tools and criteria traditionally used for assessing rCF vaginal conditions must be thoughtfully adapted for TM. This adaptation requires a fundamental shift in understanding that the microbiota considered optimal for rCF may not align with what constitutes an optimal microbiota for TM. Such redefinition is critical to developing evidence-based, individualized care strategies that address the unique needs of TM individuals.

This study provides valuable insights into the vaginal microbiota and immunological dynamics in TM individuals accessing testosterone therapy, but several limitations must be acknowledged. Firstly, the study cohort was not demographically diverse, as a significant proportion of the TM participants identified their ethno-racial identity as white or Caucasian (70.9%), and all mixed-race individuals reported Caucasian or white as part of their identity. This limitation prevents the generalizability of the findings to more diverse populations. Secondly, the data on symptoms were self-reported by participants during weekly REDCap surveys, introducing the potential for self-reporting bias. Symptoms may have been under- or over-reported depending on individual perceptions or recall, which could affect the robustness of the symptom-related findings. Finally, while the study aimed to characterize the effects of sustained testosterone use on vaginal microbiota composition and immunologic status, serum testosterone concentrations could not be measured. This limitation hampers our ability to assess whether variations in testosterone levels directly influence microbiota composition or immunological state. Future studies should measure serum testosterone levels, including upon initiation of testosterone therapy, to explore this potential relationship more comprehensively. Despite these limitations, the study provides a foundational understanding of the unique vaginal microbiota and immune dynamics in TM individuals, paving the way for more inclusive and targeted research in the future.

## RESOURCE AVAILABILITY

### • Lead contact

Dr. Jacques Ravel

### • Materials availability

This study did not generate new unique reagents.

### • Data and code availability

The 16S rRNA gene amplicon sequence datasets generated during this study are available at NCBI/SRA under accession number PRJNA1230205.

All original code and datasets utilized to conduct these analyses have been deposited in GitHub at github.com/ravel-lab/TransBiota_TM_16S/ and are publicly available.

Any additional information required to reanalyze the data reported in this paper is available from the lead contact upon request.

## ACKNOWLEDGEMENTS

Research reported in this publication was supported by the National Institute of Allergy and Infectious Diseases (NIAID) of the National Institutes of Health (NIH) under award R21AI157912. The Canadian Institutes for Health Research (CIHR PJT 180322), and the CIHR National Women’s Health Research Initiative (NWI 191322) additionally supported this work. JLP is supported by the Canada Research Chairs Program (CRC-2020-00175). AS is supported by a Canada Graduate Scholarship from CIHR. This research was undertaken, in part, thanks to funding from the Canada Foundation for Innovation (CFI 42343). The authors would like to thank Jason Hallarn and Greta Bauer for their contribution to establishing TransBiota. All sequencing was performed by Maryland Genomics (MDG) at the Institute for Genome Sciences, University of Maryland School of Medicine (UMSOM). We acknowledge assistance from Mike Humphrys at MDG with all aspects of sample collection to sequencing. Data from the Study of Women’s Health Across the Nation (SWAN) were used in this study; we would like to acknowledge the critical contribution of SWAN. The Study of Women’s Health Across the Nation (SWAN) has grant support from the National Institutes of Health (NIH), DHHS, through the National Institute on Aging (NIA), the National Institute of Nursing Research (NINR) and the NIH Office of Research on Women’s Health (ORWH) (Grants U01NR004061; U01AG012505, U01AG012535, U01AG012531, U01AG012539, U01AG012546, U01AG012553, U01AG012554, U01AG012495, and U19AG063720). The content of this article is solely the responsibility of the authors and does not necessarily represent the official views of the NIA, NINR, ORWH or the NIH. <u>Clinical Centers:</u> *University of Michigan, Ann Arbor – Carrie Karvonen-Gutierrez, PI 2021 – present, Siobán Harlow, PI 2011 – 2021, MaryFran Sowers, PI 1994-2011*; *Massachusetts General Hospital, Boston, MA – Sherri Ann Burnett Bowie, PI 2020 – Present; Joel Finkelstein, PI 1999 – 2020*; *Robert Neer, PI 1994 – 1999; Rush University, Rush University Medical Center, Chicago, IL – Imke Janssen, PI 2020 – Present; Howard Kravitz, PI 2009 – 2020*; *Lynda Powell, PI 1994 – 2009; University of California, Davis/Kaiser – Elaine Waetjen and Monique Hedderson, PIs 2020 – Present; Ellen Gold, PI 1994 - 2020*; *University of California, Los Angeles – Arun Karlamangla, PI 2020 – Present; Gail Greendale, PI 1994 - 2020*; *Albert Einstein College of Medicine, Bronx, NY – Carol Derby, PI 2011 – present, Rachel Wildman, PI 2010 – 2011; Nanette Santoro, PI 2004 – 2010; University of Medicine and Dentistry – New Jersey Medical School, Newark – Gerson Weiss, PI 1994 – 2004;* and the *University of Pittsburgh, Pittsburgh, PA – Rebecca Thurston, PI 2020 – Present; Karen Matthews, PI 1994 - 2020.* <u>NIH Program Office:</u> *National Institute on Aging, Bethesda, MD – Rosaly Correa-de-Araujo 2020 - present; Chhanda Dutta 2016- present; Winifred Rossi 2012–2016; Sherry Sherman 1994 – 2012; Marcia Ory 1994 – 2001; National Institute of Nursing Research, Bethesda, MD – Program Officers.* <u>Central Laboratory:</u> *University of Michigan, Ann Arbor – Daniel McConnell* (Central Ligand Assay Satellite Services). <u>Coordinating Center:</u> *University of Pittsburgh, Pittsburgh, PA – Maria Mori Brooks, PI 2012 - present; Kim Sutton-Tyrrell, PI 2001 – 2012; New England Research Institutes, Watertown, MA - Sonja McKinlay, PI 1995 – 2001.* Steering Committee: Susan Johnson, Current Chair, Chris Gallagher, Former Chair. We thank the study staff at each site and all the women who participated in SWAN.

## AUTHOR CONTRIBUTIONS

Conceptualization: J.R., J.L.P., Y.K, E.P. Methodology: B.M., P.H., D.Z., L.R. Formal analysis: B.M., H.W., P.G., J.R-V. Interpretation: B.M., J.L.P. and J.R. Writing: B.M., J.R., Editing: B.M., H.W., P.H., P.G., J.R-V., A.S., R.P., E.W., Y.K, E.P., J.L.P. and J.R. Funding acquisition: J.R. and J.L.P.

## METHODS

### Sample collection and study design

TM individuals who had not undergone vaginectomy and who had been on testosterone therapy for at least one year were recruited for the TransBiota study. Recruitment occurred between March 2021 and November 2022 via social media platforms and posters displayed at clinical partner sites. Eligibility criteria included absence of pregnancy and refraining from vaginal sex or the use of vaginal hygiene products for at least three days prior to sample collection.

Once a week for three weeks, participants were directed to self-collect three mid-vaginal swabs, using sterile flocked swabs. The first was placed directly into C2.1 solution (Qiagen), a validated buffer that stabilizes DNA and RNA at room temperature for at least two weeks.^84^ The second was placed into 500 µl of a stabilization buffer, containing a protease inhibitor (Complete Mini Protease Inhibitor Cocktail, Sigma-Aldrich), antimicrobial agent (Primocin, Invivogen) and 10% bovine serum albumin (Sigma-Aldrich), validated in-house to stabilize immune analytes at room temperature for up to 14 days. Finally, the third was rolled across a glass slide and used to measure pH by pH strip and participant matching to a provided color chart. Participants then mailed their sample back to University of Western Ontario (UWO), where microbiota and immune samples were stored at -80 C. Microbiota samples were shipped to the Institute for Genome Sciences at the University of Maryland School of Medicine on dry ice, where they were stored at -80 C until processing.

We hypothesized that transmasculine genital microbiotas might be similar to three groups: reproductive-aged cis women, uncircumcised cis men, and postmenopausal cis women. Reproductive-aged cis Canadian women were recruited as geographically similar comparators; each woman provided two vaginal samples, one for microbiota analyses and one for SMI characterization. As the often-anaerobic environment of the foreskin-covered coronal sulcus would be more similar to the microaerobic environment of the vagina than the aerobic environment of the circumcised coronal sulcus, uncircumcised cis men of any age were recruited to provide a single sample for microbiota analyses. Finally, we leverage a subset of 200 race-matched samples from the Study of Women’s Health Across the Nation, which characterized the vaginal microbiota of over 1300 postmenopausal cis women in the United States.^50^

The Institutional Review Board at University of Maryland School of Medicine and the Review Ethics Board at University of Western Ontario approved the protocol. Guidelines of both universities were followed in the conduct of clinical research.

### Genomic DNA extraction, 16S rRNA gene V3-V4 amplification and sequencing

Genomic DNA from samples stored in Qiagen C2.1 solution was isolated by the University of Maryland Medical School’s Maryland Genomics core facility, using the Qiagen MagAttract PowerMicrobiome DNA/RNA kit (Cat. No. 27500-4-EP) on a Hamilton STAR. Samples were thawed on ice, and 200 μl of sample was used as input, following the previously published and validated protocol described by Holm et al.^85^ Briefly, the V3-V4 regions of the 16S rRNA gene were amplified with a two-step protocol and dual index barcoding. Amplicons underwent pooling in equimolar concentrations and were purified prior to sequencing on either an Illumina MiSeq or NextSeq 1000. Negative and positive control samples were extracted and sequenced in parallel, as previously described.^85^

### Sequencing quality assessment/control and taxonomic assignment strategy

Sequences were demultiplexed via a dual barcode strategy, using a mapping file linking barcode to sample ID and split_libraries.py, a QIIME-dependent script.^86^ Resulting forward and reverse fastq files were then split by sample, using the QIIME-dependent script split_sequence_file_on_sample_ids.py, and primer sequences were removed via TagCleaner (v.0.16).^87^ Resulting paired end sequences were processed using dada2^87^ to identify amplicon sequence variants (ASVs) and remove chimeric sequences, following best practices described by https://benjjneb.github.io/dada2/bigdata.html. Each amplicon sequencing variant (ASV) generated by dada2 was assigned taxonomy using the RDP Naïve Bayesian Classifier^88^ trained with the SILVA 16S rRNA gene database (version 132).^89^ SpeciateIT and vSpeciateDB (version 1.0)^90^ were used to assign species to major genera, and sequence counts for ASVs assigned to the same taxonomy were summed for each sample. After removing samples with fewer than 500 total sequences (n = 38) and taxa present at less than 1e^-5^ overall abundance in the dataset (303 taxa), the final sequence count dataset featured 4,258,597 high-quality sequences, with a median of 21,236 reads per sample (IQR: 7,078-29,956), 262 unique taxa, and 208 samples from 85 individual participants; each participant contributed an average of 2.45 weekly samples (range: 1.0-3.0 swabs, IQR: 2.0-3.0 swabs). Relative abundances were calculated by dividing the summed sequence count of ASVs from each taxon by the total ASV sequence count for each sample.

### Assignment of vaginal community state types

The VALENCIA tool,^29^ described in 2020 by France et al., can be used to reliably and reproducibly assign vaginal and other microbiota samples to community state types (CSTs) commonly observed in the vaginal microbiota of reproductive-age cis women. For each sample included in the analyses, including transmasculine TransBiota samples (n = 208, N = 85), postmenopausal cis female vaginal samples (n = 200), reproductive-aged cis female vaginal samples (n = 35), and penile samples from uncircumcised cisgender men (n = 56), relative abundance tables were processed using the VALENCIA tool and the following command: python3 Valencia.py -i ∼/TMI_combined_SWAN_valencia_input.csv -o ∼/VALENCIA_TMI_SWAN_VC_PC_out -p ∼/VALENCIA_TMI_SWAN_VC_PC -r ∼/CST_centroids_012920.csv and the resulting CST assignment was saved.

### Vaginal immune milieu characterization

The following 13 immune analytes were selected for quantification based on previously published (a) associations with bacterial dysbiosis, inflammation, and HIV susceptibility in genital microenvironments (IL-1α, IL-1β, IL-6, IL-8, IP-10, TNF-α)^32–42^; (b) chemotactic activity (MIG, MIP-1α, MIP-1β, RANTES)^43^; (c) relevance to mechanical injury of the epidermal barrier (IL-1β, IL-6)^42,44^; and (d) associations with chronically inflammation (IL-10).

The concentrations of immune analytes were measured using a multiplex immunoassay (Millipore Sigma Multiplex Panel A kits measured on a Luminex MAGPIX system) in accordance with the manufacturer’s instructions. The immunoassay plate was washed with an automated plate washer (BioTek 405 TS washer) to standardize the washing steps. The results of the bioassays were quantified, and the lower limit of quantification (LLoQ) was assigned for each analyte based on the assessment of standard curves using Belysa^®^ Immunoassay Curve Fitting Software (Millipore Sigma Cat. No. 40-122). All samples that fell below the LLoQ were assigned a concentration of 0pg/ml for the purposes of analysis.

If most samples (>85%) fell below the determined LLoQ for a particular analyte, that analyte was removed from the final analysis. Analytes with >85% of samples above the LLoQ, but with (a) the majority of detectable samples very close in concentration to the LLoQ, and (b) poor CV (>20%) values for samples close to the LLoQ, were also assigned a lower limit of detection (LLoD). Samples above the LLoD (based on standard curves), but below the LLoQ (based on measurement replicability) were deemed detectable, but not quantifiable. Samples that fell below the LLoD were assigned a value of 0pg/ml, and samples that fell between the LLoD and the LLoQ were assigned the value halfway between the LLoD and the LLoQ. Only analytes with at least 30% prevalence after these transformations were included in the final analysis.

The concentration of SMI detected in transmasculine persons (n = 208) and cis women of reproductive age (n = 29) were compared via Kruskal-Wallis test, with significance set *a priori* at p <= 0.05, with pairwise comparisons determined using Dunn’s test.^91^

### Comparison of genital microbiota between transmasculine and cisgender individuals

The relative abundances of the top 50 most abundant taxa across four groups – transmasculine TransBiota participants, reproductive-aged cis women, postmenopausal cis women, and uncircumcised cis men – were stratified by group then sorted by VALENCIA or tmCST and relative abundances of *P. timonensis*, *Streptococcus*, *Corynebacterium* 1, and *G. vaginalis* (**Fig. 1**). To evaluate drivers of dissimilarity between the four groups, we utilized Principal Coordinates Analysis (PCoA) with Jensen-Shannon divergence. Jensen-Shannon dissimilarity indices were calculated using a customized *vegdist* function in R and projected in 2D space using the *cmdscale* function in the *stats* package in R. Resulting eigen values were transformed into percentages, and the top 2 principal components were plotted. The *envfit* function from the *BiodiversityR* package was utilized to fit taxa vectors onto each ordination, and taxa with r^2^ values greater or equal to 0.5 were plotted.

### tmCSTs and their associations with local immune status and self-reported symptoms

After removing all three samples from participant TMI162, who possessed a stable microbiota with over 95% relative abundance of *F. vaginae* at each weekly sampling, the relative abundances of taxa observed in TM microbiota samples were embedded in three-dimensional space using the PHATE^92^ tool and clustered with the HDBSCAN algorithm.^93^ We utilized *rstan*-based Bernoulli modeling with random effects correction for repeated measures to calculate the probability of symptom report per tmCST. The log_10_-transformed concentration of each soluble mediator of inflammation (SMI) was stratified by tmCST, the Kruskal-Wallis test was used to test for differences in distribution of SMI concentrations, and Dunn’s test with Bonferroni correction was applied to test for pairwise differences between tmCSTs. Counts and probabilities of transitioning between tmCSTs were calculated and displayed in a chord graph and a heatmap of probability. Yue-Clayton theta indices were calculated with the *designdist* function from the R package *vegan* using the equation, “sum(x*y)/(sum(x^2^)+sum(y^2^)-sum(x*y))”), to compare the stability of samples over a one-week period (from W1 to W2 or W2 to W3).^94^ Yue-Clayton indices for each transition from one tmCST to another were averaged and the range was displayed in **Supp. Table 4**.

### Identifying correlations between microbiota composition, local immune status, and self-reported symptoms

To further explore associations between microbiota composition, local immune status, and self-reported symptoms, Spearman correlations between the top 50 taxa and all immune analytes were calculated. The correlation matrix was transformed to a distance matrix using the equation: (1 - cormat)/2, then hierarchical clustering was performed using the *hclust* and *cutreeDynamic* functions to identify groups of taxa significantly correlated with each other, bacterial Taxa Clusters (TC). Differentially abundant taxa were identified using MaAsLin2, accounting for multiple samples from each participant as a random effect.

## SUPPLEMENTARY TABLES

**Table S1.** Local and systemic exposures included in the analyses.

| Systemic Exposures | Yes | No |
| --- | --- | --- |
| Current Anti-androgenic Medication Use | 3 (3.5) | 82 (96.5) |
| Probiotic Use, past 30 days | 6 (7.1) | 79 (92.9) |
| Antibiotic Use, past 30 days | 3 (3.5) | 82 (96.5) |
| Nicotine Use, past 30 days | 14 (16.5) | 71 (83.5) |
| Marijuana Use, past 30 days | 36 (42.4) | 49 (57.6) |
| Genital Exposures |  |  |
| Current Local Estradiol Use | 4 (4.7) | 81 (95.3) |
| HPV diagnosis ever | 5 (5.9) | 80 (94.1) |
| Herpes diagnosis ever | 8 (9.4) | 77 (90.6) |
| "Bottom" Gender Affirming Surgeries | 18 (21.2) | 67 (78.8) |
| Hysterectomy and Oophorectomy | 12 |  |
| Hysterectomy alone | 4 |  |
| Oophorectomy alone | 1 |  |
| Metoidioplasty | 1 |  |
| Steroid Use, past 30 days | 13 (15.3) | 72 (84.7) |
| Current IUD | 10 (11.8) | 75 (88.2) |
| Copper IUD | 3 |  |
| Progestin IUD | 7 |  |

**Table S2.** Self-reported symptoms experienced by transmasculine individuals currently or in the past 7 days.

| Self-Reported Symptoms, Past 7 days | N (%) |
| --- | --- |
| Dryness | 20 (23.5) |
| Pain during Sex | 11 (12.9) |
| Non-menstrual bleeding | 5 (5.8) |
| Discharge | 5 (5.8) |
| Cramping | 4 (4.7) |
| Itching or Burning Sensation | 3 (3.5) |
| Pain | 3 (3.5) |
| Tissue Thinning or Tearing | 3 (3.5) |
| "Not Normal" or Unpleasant Odor | 2 (2.4) |
| No symptoms | 48 (56.5) |

**Table S3.** Immune analytes characterized and detected in transmasculine individuals and reproductive-age cis women. The Kruskal-Wallis test was used to test for differences in distribution.

|  | Transmasculine Individuals |  | Reproductive-Aged Cis Women |  | KW p-value |
| --- | --- | --- | --- | --- | --- |
|  | Number of Samples Detected<br>In, n (%) | TM, median (range) | Number of Samples Detected<br>In, n (%) | CF, median (range) |  |
| IL-8 | 208 (100) | 2590.6 (3.5 - 6442.5) | 29 (100) | 2864.7 (132.3 - 7799.9) | 0.3228 |
| IL-1α | 207 (99.5) | 723.9 (6.8 - 17195.5) | 29 (100) | 1753.0 (431.2 - 19901.9) | **0.000308** |
| MIG | 205 (98.6) | 656.1 (7.5 - 10993.0) | 29 (100) | 704.2 (22.2 - 8375.3) | 0.2081 |
| IL-6 | 204 (98.1) | 25.6 (1.7 - 1494.3) | 29 (100) | 24.8 (2.2 - 1583.9) | 0.6802 |
| RANTES | 203 (97.6) | 95.1 (2.4 - 1293.6) | 29 (100) | 9.5 (3.0 - 428.9) | **9.659E-09** |
| IL-1β | 203 (97.6) | 130.1 (5.8 - 13264.7) | 29 (100) | 73.7 (10.2 - 1211.9) | 0.08018 |
| MIP-1β | 201 (96.6) | 54.7 (7.2 - 1345.8) | 29 (100) | 5.3 (5.3 - 103.0) | **1.682E-10** |
| IL-10 | 195 (93.8) | 44.6 (3.6 - 1198.1) | 19 (65.5) | 4.9 (3.6 - 45.6) | **9.772E-09** |
| TNF-α | 177 (85.1) | 26.7 (7.5 - 574.5) | 12 (41.4) | 9.2 (7.6 - 45.1) | **0.0000232** |
| IP-10 | 172 (82.7) | 56.0 (7.0 - 3723.9) | 28 (96.6) | 158.9 (3.8 - 1249.7) | **0.000258** |
| IFN-γ | 122 (58.7) | 14.8 (7.5 - 66.5) | 21 (72.4) | 12.7 (7.5 - 26.9) | 0.254 |
| MIP-1α | 117 (56.2) | 157.4 (18.2 - 1856.0) | 9 (31) | 39.7 (22.8 - 229.4) | **0.0061** |
| IL-22 | 68 (32.7) | 95.3 (65.5 - 187.3) | 10 (34.5) | 107.1 (69.1 - 167.3) | 0.1835 |

**Table S4.** Stability of transmasculine individual vaginal microbiome over one week, as measured by the Yue-Clayton theta index. Rows are the starting tmCST and columns are the ending tmCST for paired samples from the same individual. Yue-Clayton theta indices for each tmCST transition were averaged and the range displayed.

|  | 1 | 2 | 3 | 4 | 5 | 6 | 7 | 8 | 9 | 10 | 11 |
| --- | --- | --- | --- | --- | --- | --- | --- | --- | --- | --- | --- |
| 1 | 0.786<br>(0.423-0.997) | - | - | - | - | - | - | - | - | - | - |
| 2 | - | 0.615<br>(0.325-0.822) | - | 0.251<br>(0.251-0.251) | - | - | - | - | 0.113<br>(0.113-0.113) | - | - |
| 3 | 0.098<br>(0.098-0.098) | - | 0.617<br>(0.259-0.85) | 0.212<br>(0.212-0.212) | - | - | - | - | 0 (0-0) | - | - |
| 4 | - | 0.332<br>(0.227-0.465) | 0.546<br>(0.546-0.546) | 0.488<br>(0.068-0.929) | 0.262<br>(0.262-0.262) | - | - | 0.409<br>(0.409-0.409) | 0.027<br>(0.027-0.027) | 0.305<br>(0.305-0.305) | - |
| 5 | - | - | - | 0.132<br>(0.132-0.132) | 0.453<br>(0.128-0.626) | - | 0.396<br>(0.076-0.678) | 0.129<br>(0.129-0.129) | 0.103<br>(0.103-0.103) | - | - |
| 6 | - | - | - | - | 0.506<br>(0.506-0.506) | 0.583<br>(0.427-0.725) | 0.093<br>(0.093-0.093) | - | - | - | - |
| 7 | 0.528<br>(0.528-0.528) | 0.005<br>(0.005-0.005) | - | - | 0.225<br>(0.075-0.522) | - | 0.437<br>(0.039-0.856) | 0.556<br>(0.556-0.556) | 0.26<br>(0.038-0.559) | 0.438<br>(0.311-0.565) | - |
| 8 | - | - | - | - | - | - | 0.055<br>(0.02-0.089) | 0.536<br>(0.24-0.743) | 0.563<br>(0.563-0.563) | 0.189<br>(0.189-0.189) | - |
| 9 | - | - | 0.1<br>(0.1-0.1) | - | 0.041<br>(0.041-0.041) | - | 0.379<br>(0.379-0.379) | - | 0.518<br>(0.246-0.732) | 0.691<br>(0.691-0.691) | - |
| 10 | - | - | - | 0.134<br>(0.039-0.37) | - | - | 0.212<br>(0.045-0.341) | 0.489<br>(0.394-0.584) | 0.222<br>(0.222-0.222) | 0.661<br>(0.424-0.962) | - |
| 11 | - | - | - | - | - | - | - | - | - | - | 0.138<br>(0.044- |
|  |  |  |  |  |  |  |  |  |  |  | 0.232) |

